# Response of UBR-box E3 ubiquitin ligases and protein quality control pathways to perturbations in protein synthesis and skeletal muscle size

**DOI:** 10.1101/2025.07.23.666188

**Authors:** Leslie M. Baehr, Luis Gustavo Oliveira de Sousa, Craig A. Goodman, Adam P. Sharples, David S. Waddell, Sue C. Bodine, David C. Hughes

## Abstract

The N-degron pathway contributes to proteolysis by targeting N-terminal residues of destabilized proteins via E3 ligases that contain a UBR-box domain. Emerging evidence suggests the UBR-box family of E3 ubiquitin ligases (UBR1-7) are involved in the positive regulation of skeletal muscle mass. The purpose of this study was to explore the role of UBR-box E3 ubiquitin ligases under enhanced protein synthesis and skeletal muscle growth conditions. Cohorts of adult male mice were electroporated with constitutively active Akt (Akt-CA) or UBR5 RNAi constructs with a rapamycin diet intervention for 7 and 30 days, respectively. In addition, the UBR-box family was studied during the regrowth phase post nerve crush induced inactivity. Skeletal muscle growth with Akt-CA or regrowth following inactivity increased protein abundance of UBR1, UBR2, UBR4, UBR5 and UBR7. This occurred with corresponding increases in Akt-mTORC1/S6K and MAPK/p90RSK signaling and protein synthesis. The increases in UBR-box E3s, ubiquitination, and proteasomal activity occurred independently of mTORC1 activity and were associated with increases in markers related to autophagy, ER-stress, and protein quality control pathways. Finally, while UBR5 knockdown (KD) evokes atrophy, it occurs together with hyperactivation of mTORC1 and protein synthesis. In UBR5 KD muscles, we identified an increase in protein abundance for UBR2, UBR4 and UBR7, which may highlight a compensatory response to maintain proteome integrity. Future studies will seek to understand the role of UBR-box E3s towards protein quality control in skeletal muscle plasticity.

**New and Noteworthy:** Novel UBR-box E3 ubiquitin ligases are responsive to heightened protein synthesis and alterations in skeletal muscle mass and fiber size, in order to maintain proteome integrity.

## Introduction

Skeletal muscle is a highly adaptable tissue that can respond to various stimuli (e.g., inactivity, disease, and exercise) and has the ability to increase (i.e. hypertrophy) or decrease (i.e. atrophy) in size when required (1–3). The regulation of skeletal muscle mass is orchestrated by the activity of key signaling pathways that control protein breakdown and synthesis within myofibers (2). Early seminal work highlighted the critical role for the Akt-mTORC1-p70S6K1 signaling pathway in skeletal muscle mass regulation (4, 5). Since this discovery, our understanding of growth signals in skeletal muscle has evolved towards mTORC1-dependent and mTORC1-independent mechanisms, along with other processes that may contribute to hypertrophy, such as ribosome biogenesis (6–13). Genetic models, such as transgenic constitutively active Akt (Akt-CA) and TSC1 knockout mice, have expanded our knowledge of the mTORC1 signaling pathway and the regulation of muscle mass (14–19). Indeed, the transgenic Akt mice display a phenotype of increased skeletal muscle fiber size and protein synthesis, while electroporation-mediated transfection of an Akt-CA construct elicits similar effects on skeletal muscle growth (4, 19, 20). Previously published evidence from the TSC1 knockout mouse has observed mTORC1 to be an inhibitor of autophagy and regulate the ubiquitin proteasome system (UPS) (19, 21). The regulation of the UPS by mTORC1 directly remains to be determined as the UPS may be active in response to mTORC1-mediated activation of translation, where the UPS is needed to maintaining proteome integrity. However, given that the characterization of the UPS has predominantly centered on atrophy models, and only a handful of E3 ubiquitin ligases have been studied to date (e.g. MuRF1, MAFbx, and MUSA1) (22–24), very little is known about the UPS in skeletal muscle remodeling and growth. Thus, our understanding of the UPS, and other protein quality control systems (e.g. unfolded protein response, autophagy), could be expanded by characterizing new targets or exploring different contexts and scenarios where these systems are influential in tissue health, such as skeletal muscle growth and remodeling.

The UBR-box family of E3 ubiquitin ligases, which consists of HECT, RING and F-box E3 ubiquitin ligases named UBR1 through UBR7, encompasses part of the ubiquitin proteasome system, and their action on protein substrate recognition forms the N-degron pathway (25, 26). The N-degron pathway (also known as the N-end rule pathway) identifies degradation signals on destabilized proteins through the presence of ubiquitin and acetyl motifs (25, 27). As part of the N-degron pathway, the UBR-box family of E3 ubiquitin ligases identify unacetylated protein substrates for ubiquitination via the UBR-box domain.(25, 27–29). Studies have highlighted that UBR-box E3s play diverse roles in a range of cellular processes, with protein quality control and protein clearance typically being referenced (30–33). However, very little is known about this family of E3 ubiquitin ligases in skeletal muscle. In an early study from the Goldberg lab (34), various inhibitors targeting E3α (also known as UBR1) showed an ∼60% contribution of this proteolytic pathway to the turnover of endogenous proteins in mammalian skeletal muscle. More recently, studies have identified UBR4, UBR5 and UBR7 to be responsive to exercise stimuli (35–39) and UBR2 for being important in the development of cancer cachexia (40, 41). Our previous studies have identified a potential role for UBR5 in skeletal muscle growth and recovery following disuse (42). Further, UBR5 knockdown in mouse skeletal muscle perturbed Akt-mTORC1-p70S6K1 and MAPK/p90RSK signaling pathways, that coincided with a loss of skeletal muscle mass and fiber cross-sectional area (CSA) (43). Thus, it has been observed that UBR-box E3 ubiquitin ligases may be important for skeletal muscle health and the regulation of skeletal muscle mass (30, 36, 37, 42–44).

Therefore, we sought to investigate if UBR-box E3 ubiquitin ligases and protein quality control systems are responsive to genetic manipulation of Akt-mTORC1 signaling and the associated increase in protein synthesis and skeletal muscle size. In addition, we wanted to ascertain if these effects were dependent on mTORC1 activity. Lastly, given our previous observations of UBR5 in models of skeletal muscle regrowth following a period of atrophy (42), we hypothesized that UBR1, UBR2, UBR4 and UBR7 might display a similar temporal profile to UBR5 during the regrowth phase of the nerve crush injury model.

## Materials and methods

### Study approval for animal experiments

The mice used in these studies were males from the C57BL/6N strain obtained from Charles River laboratories at 5-6 months of age and used for experiments within 2 weeks of their arrival. Animals were housed in ventilated cages maintained in a room at 21 °C with 12-h light/ dark cycles and had *ad libitum* access to standard chow (LabDiet PicoLab® 5053) and water throughout the study. All animal procedures were approved by the Institutional Animal Care and Use Committee at the Oklahoma Medical Research Foundation.

### Rapamycin Diet Intervention

Male C57BL/6 mice received either a control diet (PicoLab® 5053 (LabDiet®) containing Eudragit) or a diet containing 42 mg/kg chow active encapsulated rapamycin (Rapa; Rapamycin Holdings, Inc., San Antonio, TX) which was provided ad libitum. The Eudragit in the chow serves as a true experimental control, given that the rapamycin is encapsulated by this material in order to provide effective release of the drug. For the Akt overexpression experiments, age-matched cohorts of mice were randomized into control or rapamycin diet interventions for 7 days prior to the electroporation procedure and remained on the respective diet until completion (total time equaled 14 days) (Figure 1). For the UBR5 knockdown experiments, age-matched cohorts of mice were electroporated and randomized into control or rapamycin diet interventions and remained on the diets for a total of 30 days (Figure 1). Chronic treatment of rapamycin has been observed to target mTORC1 and mTORC2 for inhibition (45). The rapamycin dose implemented has been reported to inhibit mTORC1 activity (46–48) and we confirmed mTORC1 inhibition by assessing p70S6K1 and rpS6 phosphorylation as downstream markers for mTORC1 activity. Body weight was assessed throughout the dietary interventions, and no difference was observed between control and rapamycin-treated mice over the 14 to 30 days intervention (data not shown).

**Figure 1.**
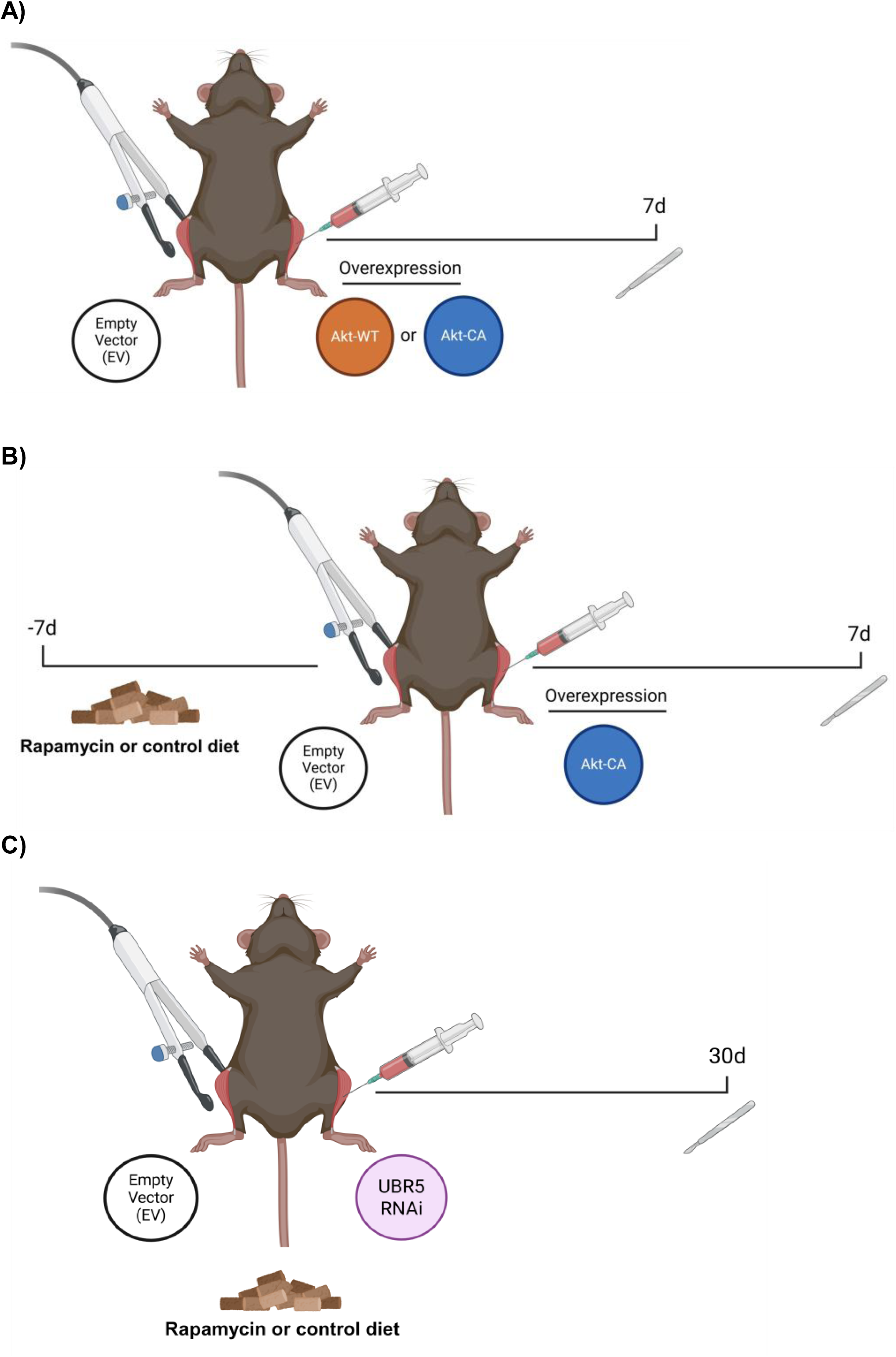
Overview of the experimental designs for studies using in vivo electroporation of the tibialis anterior (TA) muscle with dietary manipulation. Mice were electroporated with an empty vector control and either the wild type (Akt-WT) or constitutively active (Akt-CA) Akt constructs for 7 days (A). Next, cohorts of mice were randomly assigned to a control or rapamycin diet for 7 days. The mice were then electroporated with EV and Akt-CA constructs and collected after 7 days. Total time on the dietary intervention was 14 days (B). Lastly, cohorts of mice were electroporated with EV and UBR5 RNAi constructs and then placed on a control or rapamycin diet for 30 days (C). All studies utilized adult male mice (4-5 months of age). Figure was created with Biorender.com.

### Akt and UBR5 Plasmid Constructs

For the Akt experiments, the open reading frame of mouse Akt1 (Akt-WT) was cloned and ligated into the pCMV5 expression plasmid and fused with a hemagglutinin (HA) tag at the carboxyl-terminus. To create the constitutively active (Akt-CA) construct, a consensus myristylation sequence (MGSSKSKPKSR) was fused to the amino terminus of HA-tagged, wild-type mouse Akt1 in the pCMV5 expression plasmid. For RNAi experiments, the negative control RNAi plasmid was described previously (42, 49) and encodes emerald green fluorescent protein (EmGFP) and a non-targeting pre-miRNA under bicistronic control of the cytomegalovirus (CMV) promoter in the pcDNA6.2GW/EmGFP-miR plasmid (Invitrogen, Carlsbad, CA). The UBR5 RNAi plasmid encodes EmGFP and an artificial pre-miRNA targeting the full-length gene of mouse UBR5 under bicistronic control of the CMV promoter; it was generated by ligating the Mmi571982 oligonucleotide duplex (Invitrogen, Carlsbad, CA) into the pcDNA6.2GW/EmGFP-miR plasmid. The successful knockdown of UBR5 mRNA expression and protein levels in mouse skeletal muscle was previously screened and confirmed at approximately 50-60% (42, 43).

### In vivo electroporation

Transfection of mouse skeletal muscle with plasmid DNA was performed in mice under isoflurane (2-4% inhalation) anesthesia as previously described (50–52). Briefly, after a 2h pre-treatment with 0.4 units/µl of bovine placental hyaluronidase (Sigma) resuspended in sterile 0.9% saline, 20µg of plasmid DNA was injected into the tibialis anterior (TA) muscle. The hind limbs were placed between two-paddle electrodes and subjected to 10 pulses (20 msec) of 175 V/cm using an ECM-830 electroporator (BTX Harvard Apparatus, Holliston, MA). For Akt overexpression studies, an additional 2 µg of emerald-GFP plasmid was electroporated for identification of GFP positive fibers. Following transfection, mice were returned to their cages to resume normal activities until tissue collection.

### Nerve Crush Injury Model

Gastrocnemius complex (GSTC) muscle samples from our previously published study (42) were reanalyzed for protein abundance of UBR-box E3 ubiquitin ligases (UBR1, UBR2, UBR4, UBR5 and UBR7) via immunoblotting. Targeted nerve crush in the lower limb muscles of the right leg was accomplished via acute 10 s crushing of the sciatic nerve in the midthigh region of mice using forceps. The crushing of the sciatic nerve was performed by the same investigator for all cohorts of mice with firm pressure being applied for the 10 s time period. The procedure was performed under isoflurane anesthesia (3% inhalation) using aseptic surgical techniques. Male mice (C57BL/6; 12 weeks old) purchased from Charles River Laboratories (Wilmington, MA) were given an analgesic (buprenorphine, 0.1 mg kg^−1^) immediately, as well as for 48 h following surgery, and returned to their cage following recovery. Following completion of the appropriate time point (n=6/group), mice were anesthetized with 3% isoflurane, and the gastrocnemius complex (GSTC) muscles were excised, weighed, frozen in liquid nitrogen and stored at −80°C for later analysis. Prior to tissue collection, a sciatic nerve test was applied via a single electrical pulse to the sciatic nerve through electrodes to observe a twitch response in the hindlimb muscles. Between 21 and 60 days, all animals displayed a twitch, suggesting that neural activity to the hindlimb muscles had returned at these time points. A separate untreated cohort of animals (n = 4/group) was used as the relevant control. We assessed UBR-box E3 ubiquitin ligase protein abundance at time points 21-, 28-, 45-, and 60-days post nerve crush injury which coincided with reinnervation and reflective of skeletal muscle regrowth following a period of inactivity.

### Tissue Collection

Following completion of the appropriate time period following the electroporation procedure (i.e. 7 or 30 days), mice were anesthetized with isoflurane, the TA muscles were excised, weighed, frozen in liquid nitrogen, and stored at −80°C for later analysis. A subset of muscles was collected for histology and processed as described below. On completion of tissue removal, mice were euthanized by exsanguination.

### Histology

Harvested muscles were immediately fixed in 4% (w/v) paraformaldehyde for 16h at 4°C. Following a sucrose gradient incubation period (10%, 20%, and 30% for 2-3 hours each), the TA muscles were embedded in Tissue Freezing Medium (Triangle Biomedical Sciences), and a Thermo HM525 cryostat was used to prepare 10-μm serial sections from the muscle midbelly. All sections were examined and photographed using a Nikon Eclipse Ti automated inverted microscope equipped with NIS-Elements BR digital imaging software at 10x and 20x magnification for laminin and Hematoxylin & Eosin Stain respectively.

*Laminin Stain*: TA muscle sections were permeabilized in PBS with 1% triton for 10 min at room temperature. After washing with PBS, sections were blocked with 5% goat serum for 15 min at room temperature. Sections were incubated with Anti-Laminin (1:500, Sigma Aldrich, Cat no. L9393) in 5% goat serum for 2 h at room temperature, followed by two 5-min washes with PBS. Goat-anti-rabbit AlexaFluor® 555 secondary (1:333, Invitrogen Cat no. A28180) in 5% goat serum was then added for 1 h at room temperature. Slides were cover slipped using ProLong Gold Antifade reagent (Life Technologies). Image analysis was performed using Myovision software (53, 54). Skeletal muscle fiber size was analyzed by measuring ≥ 400 transfected muscle fibers per muscle (GFP-positive), per animal (10x magnification). In some muscles the transfection frequency was so high that there were few GFP-negative fibers within the same region as the GFP-positive fibers, as has been observed previously (43, 50, 51). Therefore, for Akt-CA overexpression and UBR5 knockdown (RNAi) studies, fiber size comparisons were made between the GFP-positive fibers in the EV controls and Akt-CA or UBR5 RNAi transected muscles with and without the rapamycin diet.

*Hematoxylin & Eosin Stain*: A standard H&E protocol was performed (adapted from Parlee et al. (55)). Briefly, TA muscle sections at room temperature were incubated in deionized H_2_O for 2 min on a rocker at 30 rpm (Labnet International Inc). Next, slides were incubated with Mayer’s hematoxylin solution (EMS SKU: 26043-05) for 10 min to stain the nuclei, rinsed in running H_2_O until clear, dipped 3 times in blue solution, and rinsed under running H_2_O for 2 min. This was followed by incubation in 80% ethanol for 2 min. TA sections were stained with 0.1% Eosin (Sigma E4382-25G) solution for 10 s, followed by sequential immersion in graded ethanol concentrations for 2 min each (1 x 80%, 2 x 95%, 2 x 100%). Slides were then incubated in xylene solution (three times, 3 min each). A coverslip was applied using Permount™ media (Thermo Fisher Scientific, Waltham, MA). Slides were placed horizontally at room temperature to air dry for at least 24 hours before imaging. The percentage of aberrant myofibers was calculated by dividing the number of fibers displaying aberrant/vacuole structures by the total number of fibers (19, 56, 57). An average of 1500-1800 myofibers were counted per animal (n = 4-5/group), and quantification was performed using ImageJ software (National Institutes of Health, Bethesda, MD, USA).

### Muscle Protein Synthesis (Puromycin Incorporation)

Changes in muscle protein synthesis (MPS) were assessed in TA muscles transfected for 7 and 30 days by measuring the incorporation of exogenous puromycin into nascent peptides as described previously (6, 58). Puromycin (EMD Millipore, Billerica, MA; Cat. No. 540222) was dissolved in sterile saline and delivered (0.02 μmol/g body wt by ip injection) 30 min before muscle collection. Protein synthesis was measured under fed conditions and studied in the light cycle.

### Immunoblotting

Frozen TA muscles were homogenized in sucrose lysis buffer (50 mM Tris pH 7.5, 250 mM sucrose, 1 mM EDTA, 1 mM EGTA, 1% Triton X 100, 50 mM NaF). The supernatant was collected following centrifugation at 8,000 *g* for 10 min and protein concentrations were determined using the 660 protein assay (Thermo Fisher Scientific, Waltham, MA). Twenty micrograms of protein were subjected to SDS-PAGE on 4-20% Criterion TGX stain-free gels (Bio-Rad, Hercules, CA) and transferred to polyvinylidene diflouride membranes (PVDF, Millipore, Burlington, MA). Membranes were blocked in 3% nonfat milk in Tris-buffered saline with 0.1% Tween-20 added for one hour and then probed with primary antibody overnight at 4°C. Membranes were washed and incubated with HRP-conjugated secondary antibodies at 1:10,000 for one hour at room temperature (Cell Signaling technology Cat no. 7076 and Cat no. 7074). Immobilon Western Chemiluminescent HRP substrate was then applied to the membranes prior to image acquisition. Image acquisition and band quantification were performed using the ChemiDoc MP System and Image Laboratory 6.1 software (Bio-Rad), respectively. Total protein loading of the membranes captured from images using stain-free gel technology was used as the normalization control for all blots. The following antibodies were used at 1:1000 concentration unless otherwise stated: Total Ubiquitin (Millipore, RRID:AB_2043482 and Cell Signaling, RRID:AB_2799235), UBR1 (ProteinTech, Cat no. 260069), UBR2 (Abcam, Cat no. 217069), UBR4 (Abcam, Cat no. 86738), UBR5 (Protein Tech, Cet no. 66937), UBR7 (Novus Biologicals, Cat no. NBP1-88409), VCP (Protein Tech, RRID:AB_2214635), p62 (Sigma, RRID:AB_1841064), LC3B (Sigma, Cat no. L7543), eIF2A (Protein Tech, RRID:AB_2096489), NDRG1 (Protein Tech, RRID:AB_2880676). Cell Signaling Technologies (Danvers, MA) – K48 Ub-linkage (RRID:AB_10859893), phospho-p44/42 MAPK^Thr202/Tyr204^ (RRID:AB_2315112), phospho-p90RSK^Ser380^ (RRID:AB_2687613), p90RSK (RRID:AB_659900), phospho-Akt^Ser473^ (RRID:AB_2315049), phospho-Akt^Thr308^ (RRID:AB_2629447), Akt (RRID:AB_329827), phospho-NDRG1^Thr346^ (RRID:AB_10693451), Raptor (RRID:AB_561245), phospho-p70S6K Thr^389^ (RRID:AB_330944), p70S6K (RRID:AB_331676), phospho-rpS6^Ser240/244^ (RRID:AB_10694233), rpS6 (RRID:AB_331355), phospho-4EBP1 ^Thr37/46^ (RRID: AB_560835), 4EBP1 (RRID:AB_2097841), eIF4E (RRID:AB_823488), eIF4G (RRID:AB_2096025), BiP (RRID:AB_2119845), PDI (RRID:AB_2156433), CHOP (RRID:AB_2089254). EMD Millipore–puromycin (RRID:AB_2566826). Anti-rabbit IgG, HRP-linked (1:10 000, Cell Signaling, Cat no. 7074) and anti-mouse IgG, HRP-linked (1:10 000, Cell Signaling, Cat no. 7076) were used as secondary antibodies.

### Proteasome Activity Assays

20S and 26S proteasome activities were performed as previously described (59, 60). The 26S ATP-dependent assays were performed in homogenization buffer (50 mM Tris, 150 mM NaCl, 1 mM EDTA, 5 mM MgCl_2_ [pH 7.5] and 0.5mM dithiothreitol (DTT)) with the addition of 100 μM ATP. The 20S ATP-independent assays were carried out in assay buffer containing 25 mM HEPES, 0.5 mM EDTA, and 0.001% SDS (pH 7.5). Proteasome activities were determined by adding substrates at 100 µM: Z-Leu-Leu-Glu-MCA (Peptide Institute, catalog no. 3179-v), Boc-Leu-Ser-Thr-Arg-AMC (Bachem, catalog no. I-1940), or succinyl-Leu-Leu-Val-Tyr-7-AMC (Bachem, catalog no. I-1395), for β_1_-(caspase-like), β_2_-(trypsin-like), and β_5_-subunits (chymotrypsin-like), respectively. Each assay was conducted in the absence and presence of the proteasome inhibitor bortezomib (Cell Signaling, catalog no. 2204) at a final concentration of 2 mM (β_5_) or 10 mM (β_1_ and β_2_). All proteasome assays were conducted using 10 µg of protein/well on a 96-well plate (Greiner Bio-One, catalog no. 655076), and each sample was loaded in triplicate on the plate. The activity of the 20S and 26S proteasome was measured by calculating the difference between fluorescence units recorded with or without the inhibitor in the reaction medium. Fluorescence was measured using a Spectra Max M2 Fluorescent Microplate reader (Molecular Devices, Sunnyvale, CA; excitation wavelength, 360 nm; emission wavelength, 460 nm) at 15-min intervals for 2 h. To compare all groups on the same plate e.g. control vs rapamycin diet and empty vector vs Akt-CA constructs, we utilized a sample of N=4 per group to allow for technical replicates on the same assay plate.

### Gene expression by Quantitative RT-PCR in skeletal muscle

Frozen muscle powder was homogenized using RNAzol RT reagent (Sigma-Aldrich, St Louis, MO) in accordance with the manufacturer’s instructions. cDNA was synthesized using a reverse transcription kit (High-Capacity cDNA synthesis kit; Applied Biosystems, Waltham, MA) from 1 µg of total RNA. PCR reactions (10 µL) were set up as: 2 µL of cDNA, 0.5 µL (10 µM stock) forward and reverse primer, 5 µL of Power SYBR Green master mix (Thermo Fisher Scientific, Waltham, MA) and 2 µL of RNA/DNA free water. Gene expression analysis was then performed by quantitative PCR on a Quantstudio 6 Flex Real-time PCR System (Applied Biosystems, Waltham, MA) using the mouse primers shown in Table 1. PCR cycling comprised: hold at 50°C for 2 min, 10 min hold at 95°C, before 40 PCR cycles of 95°C for 15 s followed by annealing temp (see Table 1) for 30 s, and extension at 72°C for 30 s). Melt curve analysis at the end of the PCR cycling protocol yielded a single peak. As a result of reference gene instability, gene expression was normalized to tissue weight and subsequently reported as the fold change relative to empty vector control muscles, as described previously (61). This type of analysis has previously been used extensively by our group (62–64).

### Statistical Analysis

Results are presented as mean ± standard error of measure (SEM) for all experiments. Statistical differences for in vivo overexpression studies were determined using a paired students’ t-test on muscle mass and fiber CSA data sets. For data sets with the Akt-Wt and Akt-CA construct comparisons, a one-way analysis of variance (ANOVA) was performed followed by Dunnett’s test for multiple comparisons. For data sets generated with diet (control and rapa) and construct experiments, a two-way ANOVA was performed. When applicable, a Šídák’s post hoc test was conducted for multiple comparisons across groups. Potential outliers within data sets were identified and excluded using a Grubbs test. All statistical analyses were performed using GraphPad Prism (GraphPad Software, Inc., La Jolla, CA). Results were considered significant when P ≤ 0.05.

## Results

### Effect of differing levels of Akt activation on mTORC1 signaling and skeletal muscle mass and fiber cross-sectional area

To explore the role of protein quality control systems in response to skeletal muscle growth, we utilized an Akt overexpression model where either a wild-type (WT) or constitutively active (CA) Akt construct was electroporated into male TA skeletal muscles for 7 days (Figure 1A). We observed a significant increase in TA muscle mass with the Akt-WT (+4%) and Akt-CA (+12%) constructs when compared to the respective empty vector (EV) control (Figure 2A; P ≤ 0.05). Skeletal muscle fiber CSA was assessed in EV control and Akt-CA transfected TA muscles with GFP-positive fibers being analyzed (Figure 2B). We observed a shift in GFP-positive fiber CSA distribution towards both smaller and larger fiber sizes with the Akt-CA construct verses the EV control (Figure 2B). Lastly, overexpression of Akt-WT and Akt-CA increased Akt phosphorylation and phosphorylation of downstream mTORC1 substrates, rpS6 and p70S6K1, compared to the EV control (Figure 2C and 2E). In addition, we saw a significant increase in protein abundance for eIF4E, eIF4G and puromycin labeling, indicating elevated protein synthesis with Akt-CA overexpression (Figure 2C and 2E).

**Figure 2.**
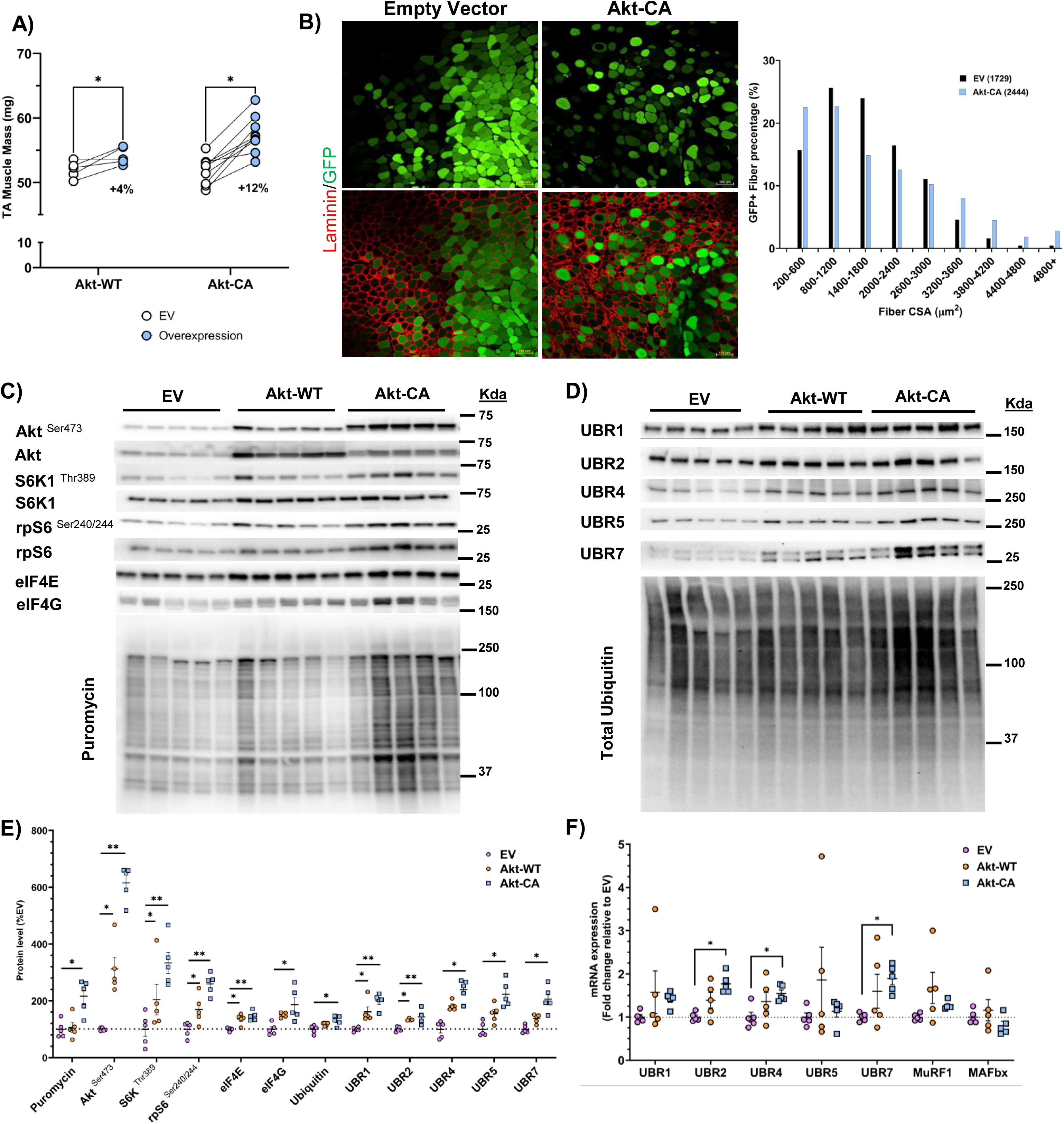
Differing degrees of Akt activity result in skeletal muscle growth and activation of mTORC1 signaling, translation and increased UBR-box E3 ubiquitin ligase abundance. Muscle mass (**A**) measurements (milligrams) for muscles transfected with either the empty vector (EV) or Akt-WT, and Akt-CA plasmids after 7 days (n=5-10/group). Representative microscope images (**B**) for green fluorescent protein (GFP) and laminin (red) in TA muscles transfected with EV or Akt-CA constructs (x10 magnification; scale bar = 100µm), highlighting transfection efficiency and muscle morphology after 7 days. Muscle cross-sectional area (CSA) distribution for EV and Akt-CA transfected muscles. GFP-positive fibers were measured for CSA, with ≥ 450 transfected fibers analyzed per animal, per muscle (n=5/group). Total number of fibers analyzed per group are reported in parentheses. For CSA data, fibers presented as percentage of fibers between 0 and 4800 µm plus size. Representative immunoblot images (**C**) for Akt/mTORC1 signaling and puromycin. Representative immunoblot images (**D**) for UBR-box E3 ubiquitin ligases and total ubiquitin. Quantification of Akt/mTORC1 signaling, puromycin-labelling, UBR-box E3 ubiquitin ligases (UBR1, UBR2, UBR4, UBR5 and UBR7) and total ubiquitin (**E**) in EV, Akt-WT, and Akt-CA transfected muscles (n=5/group). Total protein loading was used as the normalization control for all blots. In EV, Akt-WT, and Akt-CA transfected muscles, mRNA expression (**F**) was determined via quantitative PCR for UBR1, UBR2, UBR4, UBR5, UBR7, MuRF1 (Trim63), and MAFbx (FBXO32). N=4-5/group. Data presented as means ± SEM. *P<0.05; **P<0.01.

### Changes in UBR-box E3 ubiquitin ligase protein abundance and mRNA expression in Akt-transfected skeletal muscles

Given the proposed role for UBR-box E3 ubiquitin ligases in protein quality control and previous literature observing UBR5 to be elevated in models of skeletal muscle growth (35, 42), we assessed UBR-box E3 ubiquitin ligase expression in experimental models where protein synthesis and skeletal muscle mass are increased. In our Akt-WT and Akt-CA overexpression models, we observed increases in the abundance of UBR1, UBR2, UBR4, UBR5 and UBR7 proteins (Figure 2D and 2E). In addition, we observed increased levels of protein ubiquitination only in the Akt-CA transfected muscles compared to the EV control (Figure 2D). At the mRNA level, we found increased expression of UBR2, UBR4 and UBR7 in Akt-CA transfected muscles (Figure 2F). We observed no changes in UBR1, UBR5, MuRF1 or MAFbx mRNA expression with either Akt-WT or Akt-CA compared to the EV control (Figure 2F).

### UBR-box E3 ubiquitin ligases are elevated during a period of skeletal muscle regrowth following neural inactivity

To extend our observations, we analyzed UBR-box E3 ubiquitin ligase protein abundance in gastrocnemius muscle from a previously published nerve crush injury model (42). Between 21 and 60 days post nerve crush injury, skeletal muscle regrowth occurs following a period of neural inactivity (42). All the UBR-box E3 ubiquitin ligases displayed increased protein abundance between 21 and 45 days post-surgery (Figure 3A and 3B). Only UBR1 and UBR7 protein remained elevated above control tissue levels at the 60 day time point (Figure 3B). The changes in protein abundance for UBR1, UBR2, UBR4 and UBR7 correspond with our previously published observations for increased UBR5 protein abundance in the nerve crush injury model (42) during a reinnervation period when skeletal muscle mass and the protein synthetic machinery are remodeling.

**Figure 3.**
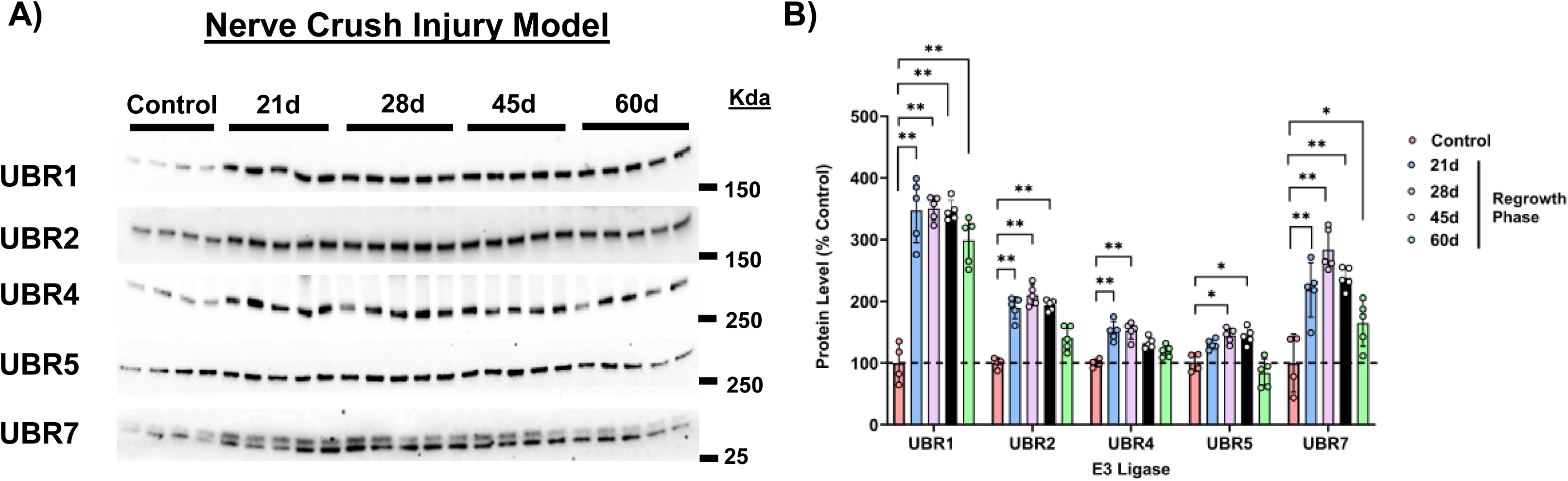
Response of UBR-box E3 ligases to skeletal muscle regrowth following a period of neural inactivity. Representative immunoblot images (**A**) for UBR-box E3 ubiquitin ligases post nerve crush injury in gastrocnemius complex (GSTC) skeletal muscle. We have previously published the UBR5 protein response in the GSTC muscle following nerve crush injury (NCI) and the 21 to 60-day time period corresponds to the recovery (regrowth) of skeletal muscle mass following a period of disuse. (**B**) Quantification of UBR-box E3 ubiquitin ligases (UBR1, UBR2, UBR4, UBR5, and UBR7) in GSTC muscle from control and time points (21, 28, 45, and 60 days post NCI) corresponding to skeletal muscle regrowth (n=4-5/group). Data presented as means ± SEM. *P<0.05, **P<0.01 vs control.

### Effect of rapamycin on Akt-induced skeletal muscle growth, rapamycin-sensitive and insensitive signaling

Next, we investigated whether changes in the content of these ubiquitin ligases with Akt overexpression were dependent on mTORC1 activity (Figure 1B), through the implementation of a rapamycin diet. In the control diet, there was a significant increase in TA muscle mass with Akt-CA overexpression, whereas the rapamycin diet blunted Akt-CA induced growth (Figure 4A). At the fiber CSA level, GFP-positive fibers were distributed towards larger and smaller size compared to the EV transfected muscles under the control diet with Akt overexpression (Figure 4B and 4C). However, the GFP-positive fibers in the rapamycin diet condition displayed only smaller fiber size in the Akt-CA transfected muscles, with a reduction in the percentage of larger fibers being observed (Figure 4B and 4C).

**Figure 4.**
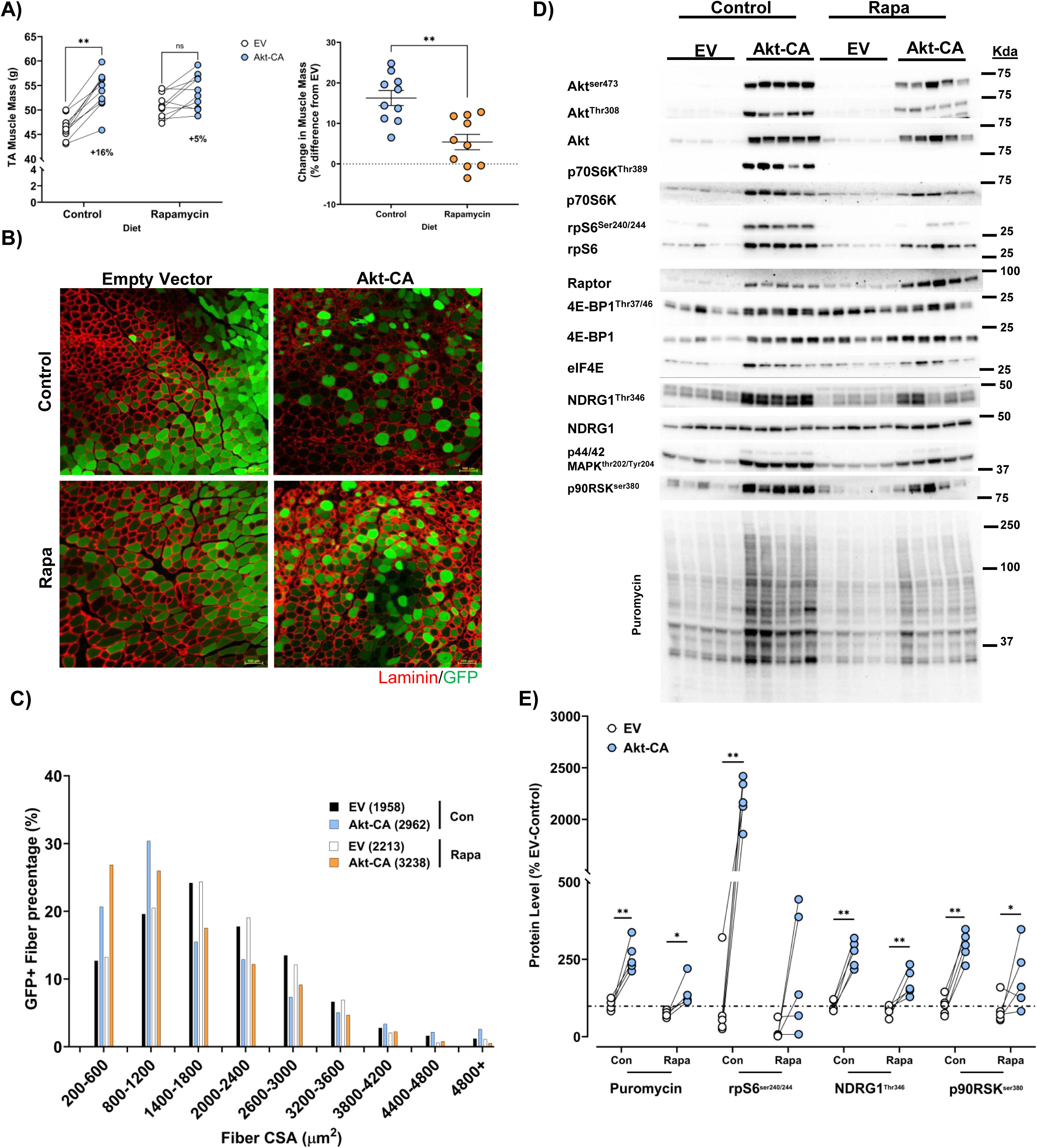
Effect of rapamycin on Akt-induced skeletal muscle growth, mTORC1 dependent and independent signaling. Male mice were provided either a control or rapamycin diet seven days prior to the electroporation procedure and kept on the respective diet for the remainder of the experiment. Muscle mass (**A**) measurements (milligrams) for muscles transfected with either the empty vector (EV) or Akt-CA plasmids for the control and rapamycin interventions after 7 days (n=10/group; left panel). The percentage difference in muscle mass compared to the EV transfected muscle for the control and rapamycin diets (A; right panel). Representative microscope images (**B**) for green fluorescent protein (GFP) and Laminin (red) in TA muscles transfected with EV or Akt-CA plasmid on the control or rapamycin diet (x10 magnification; scale bar = 100µm), highlighting transfection efficiency and muscle morphology after 7 days. Muscle cross-sectional area (CSA) distribution for EV and Akt-CA transfected muscles on the control or rapamycin diet (**C**). GFP-positive fibers were measured for CSA, with ≥ 450 transfected fibers analyzed per animal, per muscle (n=5/group). Total number of fibers analyzed per group are reported in parentheses. For CSA data, fibers presented as percentage of fibers between 0 and 4800 µm plus size. Representative immunoblot images (**D**) for Akt/mTORC1 signaling and puromycin from EV and Akt-CA plasmids with control or rapamycin diet (n=5/group). Quantification of puromycin and phosphorylation status of rpS6, NDRG1 and p90RSK in control and rapamycin conditions with Akt overexpression (**E**). Data presented as means ± SEM. *P<0.05; **P<0.01.

In terms of Akt-mTORC1 signaling, we observed increased phosphorylation of Akt, S6K and rpS6 under control diet conditions with Akt-CA overexpression (Figure 4D and 4E). Conversely, these downstream targets of mTORC1 were either non-detectable or blunted in the rapamycin diet condition (Figure 4D). However, rapamycin did not impact 4E-BP1 phosphorylation, suggesting that mTORC1 signals might not have been fully blocked. Further, we report increased protein abundance for raptor and eIF4E under control and rapamycin diet conditions with Akt-CA overexpression. Given that Akt is upstream of mTORC1 activation and can initiate signaling of other independent pathways, we assessed mTORC2 and p44/42 MAPK activity. Phosphorylation of NDRG1 (mTORC2 substrate), p44/42 MAPK and p90RSK were all significantly elevated irrespective of the diet conditions in transfected TA muscles with Akt-CA (Figure 4E). Lastly, puromycin-labeling for nascent peptides was increased in Akt-CA overexpression muscles irrespective of mTORC1 activity, highlighting the contribution of rapamycin-insensitive mechanisms to increases in the protein synthetic machinery (Figure 4D and 4E). It should be noted that the magnitude of increase for the puromycin-labelling was greater in the control group verses the rapamycin treated groups.

### Changes in UBR-box E3 ubiquitin ligases, ubiquitination and proteasomal activity are independent of mTORC1 activity

In control and rapamycin diet conditions, we assessed UBR-box E3 ubiquitin ligase protein levels to ascertain if changes occurred in a rapamycin-sensitive or insensitive manner. Protein levels for UBR2, UBR4, UBR5 and UBR7 significantly increased in the Akt-CA overexpression muscles independent of the diet condition (Figure 5A and 5B). Interestingly, UBR1 increased in the Akt-CA overexpression muscles, but also appeared to be responsive to rapamycin treatment as we observed elevated levels of UBR1 in the EV control plus rapamycin condition (Figure 5B). To further assess the UPS, we measured total ubiquitinated proteins, K48-specific linkage levels and proteasome activity for 20S and 26S subunits (Figure 5A, 5C and 5D). Independent of diet condition, we observed significantly increased levels of total ubiquitinated proteins and K48-specific linkage in Akt-CA overexpression muscles compared to the EV control (Figure 5C). For the 20S subunit proteasomal activity, we observed little change in activity with Akt-CA overexpression, except for 20S β1 activity under the rapamycin diet background (Figure 5D). In comparison, 26S proteasomal activity displayed significant elevations in β1 and β2 subunit activity in Akt-CA overexpression muscles independent of the diet condition (Figure 5D). For the 26S β5 subunit, we observed a near significant increase in activity with the Akt-CA plus rapamycin condition compared to the control diet background (Figure 5D; P = 0.051). Altogether, these data highlight changes in UBR-box E3 ubiquitin ligases, ubiquitination, and proteasomal activity that occur in a condition of low mTORC1 activity.

**Figure 5.**
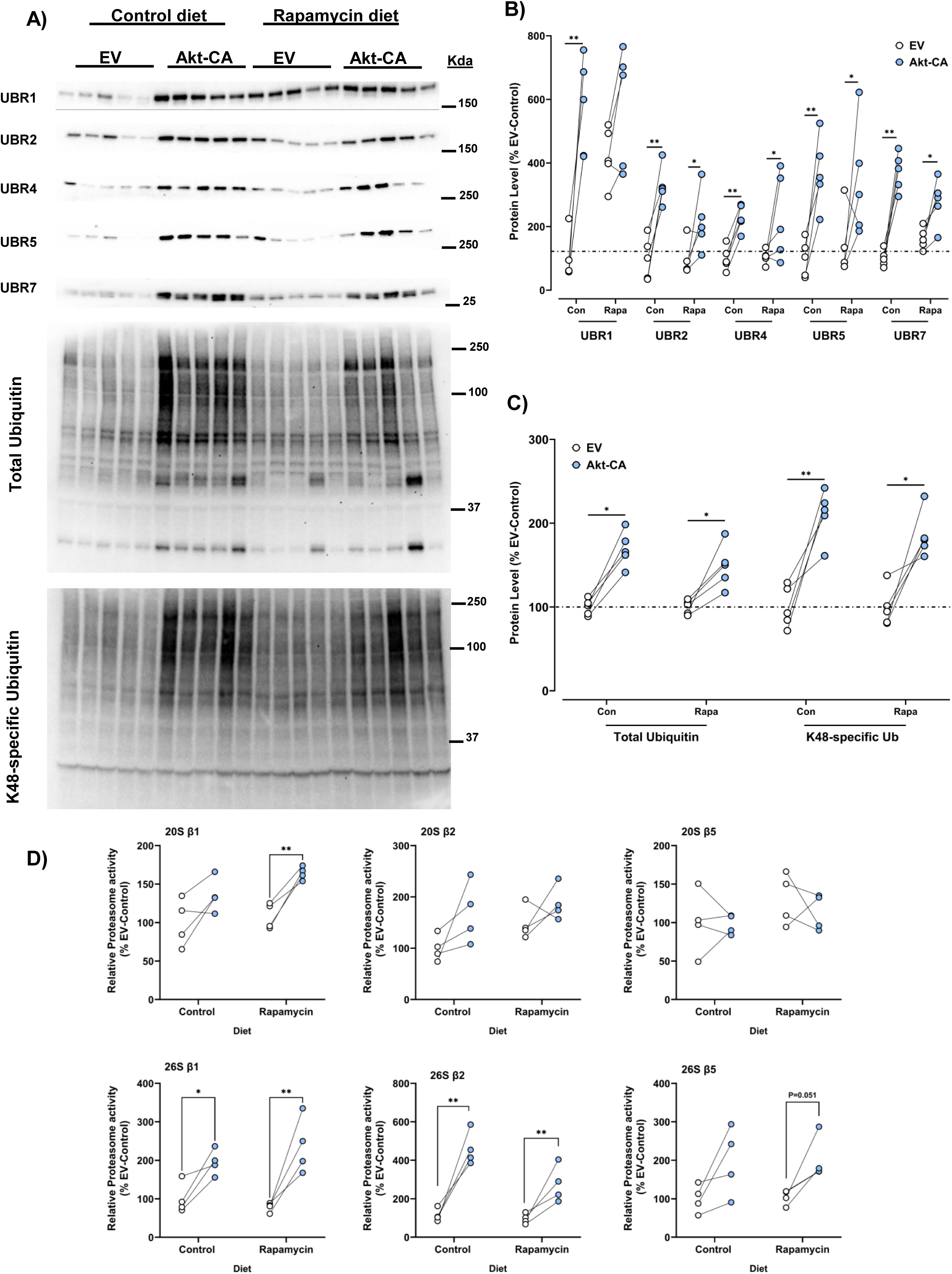
Increase in UBR-box E3 ligase abundance and proteasome activity in Akt transfected skeletal muscles under suppressed mTORC1 activity. Representative immunoblot images (**A**) for UBR-box E3 ubiquitin ligases, total ubiquitin, and K48-specific ubiquitin linkage. Quantification of UBR-box E3 ubiquitin ligases (UBR1, UBR2, UBR4, UBR5, and UBR7) (**B**), total ubiquitin, and K48 specific ubiquitin (**C**) in empty vector (EV) and Akt-CA transfected muscles under control or rapamycin diet conditions (n=5/group). Total protein loading was used as the normalization control for all blots. Proteasome activity assays were performed for 20S and 26S subunits (β1, β2, and β5) in transfected muscles (n=4/group; **D**). Data presented as means ± SEM. *P<0.05; **P<0.01.

### Markers for autophagy and ER-stress are elevated in Akt transfected muscles with suppressed mTORC1 activity

Given the reported association between the UBR-box E3 ubiquitin ligases and their involvement in protein quality control (27, 30, 31), we sought to assess changes in markers related to autophagy and ER-stress pathways in Akt-CA overexpression TA muscles. For VCP, LC3B I and LC3B II, we observed significant increased protein levels in Akt-CA overexpression muscles that were independent of diet background (Figure 6A and 6B). Interestingly for p62 protein abundance we observed significant increases in the control diet condition only with Akt-CA overexpression (Figure 6B). However, similar to UBR1, we observed a rapamycin-induced increase in basal p62 protein levels in the EV control muscles (Figure 6B). For markers related to the ER stress response, there was no change in protein levels for BiP (Figure 6C and 6D). However, for PDI, CHOP and eIF2α protein levels, there was a significant increase in Akt-CA transfected muscles that was present independent of the diet conditions (Figure 6D). Lastly, similar to Akt transgenic and TSC1/2 knockout rodent models (19), we observed the presence of ‘aberrant’ muscle fibers containing multiple vacuole-like structures in the Akt-CA overexpression muscles, which were also present in the rapamycin diet condition (Figure 6E and 6F). Overall, these data highlight the response of protein quality control pathways to Akt induction, and that these pathways may be responding to protein clearance and removal of damaged/misfolded proteins that are driven by hyperactive protein synthesis, leading to potential increased errors of translation.

**Figure 6.**
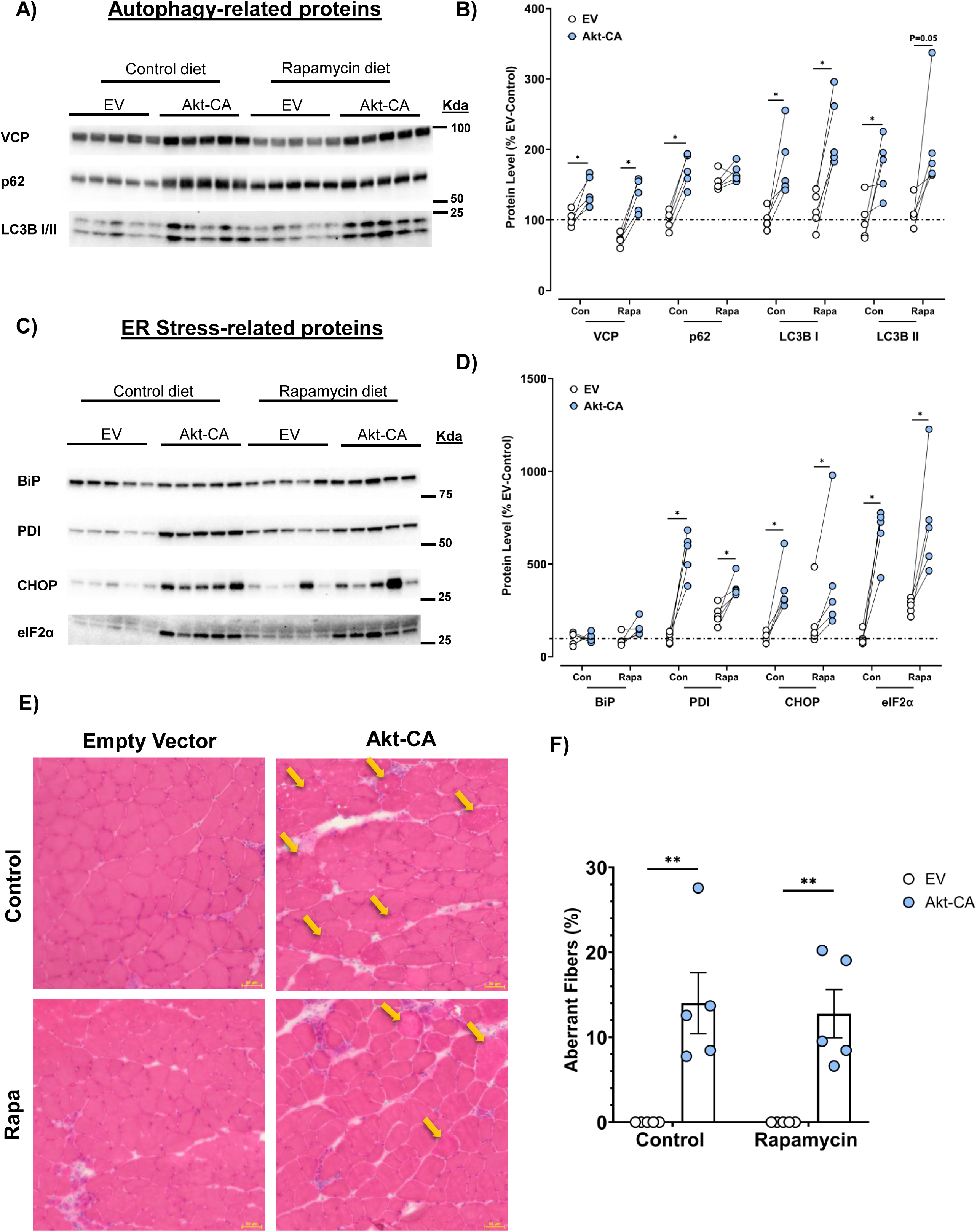
Changes in autophagy-related and ER stress-related proteins with Akt transfected skeletal muscle independent of mTORC1 activity. Representative immunoblot images (**A**) for autophagy-related markers, VCP, p62, and LC3B. Quantification of autophagy-related markers (**B**) in empty vector (EV) and Akt-CA transfected muscles under control or rapamycin diet conditions (n=5/group). Representative immunoblot images (**C**) for ER stress-related markers, BiP, PDI, CHOP, and eIF2α. Quantification of ER stress-related markers (**D**) in EV and Akt-CA transfected muscles under control or rapamycin diet conditions (n=5/group). Total protein loading was used as the normalization control for all blots. Representative microscope images for hematoxylin-eosin (H&E) in TA mouse muscles transfected with EV and Akt-CA plasmids under control and rapamycin diet conditions (×20 magnification; scale barL=L50 µm) highlighting muscle morphology after 7 days post electroporation (**E**). Examples of aberrant skeletal muscle fibers are identified on H&E images with yellow arrows and were quantified in EV and Akt-CA transfected muscles under control and rapamycin conditions (*n*=5/group) (**F**). Data presented as means ± SEM. *P<0.05; **P<0.01.

### UBR5 knockdown results in loss of muscle mass and fiber size under control and rapamycin conditions

We have previously observed UBR5 knockdown to result in loss of muscle mass and fiber size which was accompanied by mTORC1 hyperactivation, as measured by phosphorylation of downstream substrates p70S6K and rpS6 at multiple time points (43). However, it remained to be determined if the response of mTORC1 activity was adaptive or maladaptive and could be contributing to the development of skeletal muscle atrophy with UBR5 knockdown. Therefore, we sought to utilize the rapamycin diet in conjunction with UBR5 knockdown for 30 days post electroporation (Figure 1C). In the control diet group, we observed significant loss of skeletal muscle mass and fiber size, as measured by a reduction in muscle weight and a shift toward smaller CSA size for GFP-positive fibers for the UBR5 RNAi construct compared to the contralateral EV muscle (Figure 7A-C). These observations confirmed our previous findings for the effect of UBR5 knockdown on skeletal muscle mass size (43). Interestingly, we also observed significant reductions in muscle mass and fiber CSA size in UBR5 RNAi transfected muscles compared to the contralateral EV in the presence of rapamycin (Figure 7B and 7C).

**Figure 7.**
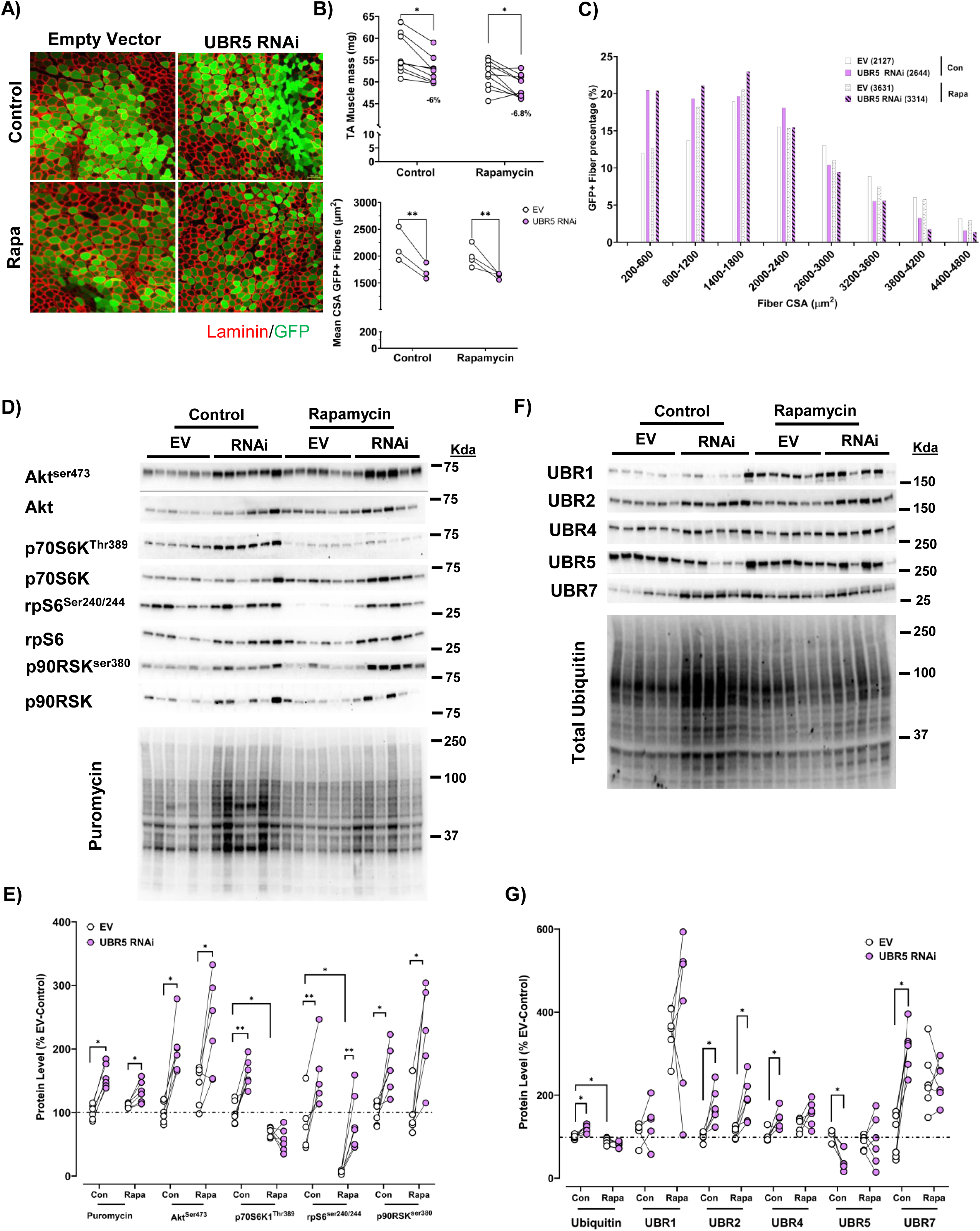
Loss of skeletal muscle mass and fiber size with UBR5 knockdown occurs independent of mTORC1 activity. Representative microscope images (**A**) for green fluorescent protein (GFP) and Laminin (red) in TA muscles transfected with EV or UBR5 RNAi plasmid on the control or rapamycin diet (x10 magnification; scale bar = 100µm), highlighting transfection efficiency and muscle morphology after 30 days. TA muscle mass (**B**) measurements (milligrams) for muscles transfected with either the EV or UBR5 RNAi plasmids for the control and rapamycin interventions after 30 days (n=9-10/group). Mean GFP-positive fiber size (**B**) was quantified for TA muscles transfected with EV or UBR5 RNAi under control or rapamycin diets (n=3-4/group). Muscle fiber CSA distribution for EV and UBR5 RNAi transfected muscles (**C**). GFP-positive fibers were measured for CSA, with ≥ 450 transfected fibers analyzed per animal, per muscle (n=3-4/group). Total number of fibers analyzed per group are reported in parentheses and represent Fiber CSA data for TA transfected muscles. For CSA data, fibers presented as percentage of fibers between 0 and 4800 µm. Representative immunoblot images (**D**) for Akt/mTORC1 signaling and puromycin from EV and UBR5 RNAi plasmids with control or rapamycin diet (n=5-6/group). Quantification of puromycin and phosphorylation status of Akt, p70S6K1, rpS6, and p90RSK in control and rapamycin conditions with UBR5 knockdown (**E**). Representative immunoblot images (**F**) for UBR-box E3 ubiquitin ligases. Quantification of UBR-box E3 ubiquitin ligases (UBR1, UBR2, UBR4, UBR5, and UBR7) and total ubiquitin (**G**) in EV and UBR5 RNAi transfected muscles (n=5-6/group) under control and rapamycin diet conditions. *P<0.05 vs. EV. Total protein loading was used as the normalization control for all blots. Data presented as means ± SEM. *P<0.05; **P<0.01.

At the biochemical level, similar to our previously published data (43), we observed increased phosphorylation of Akt, p70S6K1, rpS6 and p90RSK in UBR5 RNAi transfected muscles in the control diet condition compared to the EV muscles (Figure 7D). The phosphorylation of p70S6K at Thr389 was suppressed in the EV and UBR5 RNAi transfected muscles confirming the inhibitory effect of the rapamycin diet on mTORC1 activity (Figure 7D and 7E). Interestingly, rapamycin did not inhibit the UBR5 RNAi-induced phosphorylation of Akt, rpS6 and p90RSK sites compared to the EV transfected muscle (Figure 7D and 7E). The dissociation between p70S6K1 and rpS6 phosphorylation is interesting and might suggest the potential involvement of either another rpS6 kinase (e.g. PKA) or the inhibition of a rpS6 phosphatase (65–67). This observation warrants further investigation in the UBR5 knockdown model. Lastly, we wanted to observe how the protein abundance for UBR1, UBR2, UBR4, and UBR7 might change with UBR5 Knockdown. In the control diet, we observed significant increases in total ubiquitin, UBR2, UBR4, and UBR7 when UBR5 protein was suppressed 50-60% (Figure 7F and 7G) In the rapamycin diet condition, basal UBR1 and UBR7 proteins were significantly increased in EV transfected muscles compared to control diet EV transfected muscles, while only UBR2 remained significantly elevated in the UBR5 RNAi tissues compared to the EV transfected muscle (Figure 7F and 7G).

## Discussion

Skeletal muscle can regulate its size in response to internal and external cues throughout the life span (2, 3, 68). Homeostasis of skeletal muscle size occurs through a balance between protein synthesis and breakdown (2). In the present study, we sought to further our understanding of skeletal muscle proteostasis under growth and remodeling conditions by using a short term Akt overexpression model, in conjunction with a rapamycin diet intervention. Our results highlight a novel group of E3 ubiquitin ligases (UBR1, UBR2, UBR4, UBR5, and UBR7) that appear to be responsive under conditions of heightened protein synthesis and when other protein quality control pathways are active. In addition, the change in protein abundance for UBR-box E3 ubiquitin ligases was still evident with rapamycin treatment, suggesting that the UBR-box E3 ubiquitin ligases might act in a rapamycin-insensitive manner and maybe dependent on upstream Akt signals and mTORC1-independent control. Indeed, we observed the activation of p44/p42 MAPK, p90RSK and mTORC2 signaling that can contribute to protein synthesis and skeletal muscle growth (69–73). Future studies are required to ascertain a physiological role of UBR-box E3 ubiquitin ligases with Akt-mediated protein synthesis and turnover in the context of skeletal muscle growth and remodeling.

As the UBR-box E3 ubiquitin ligases are part of the N-degron pathway, they contain a UBR-domain which can recognize destabilized proteins and protein fragments for protein degradation (25, 27). The N-degron pathway serves as an important protein quality control mechanism (25, 31) and limited studies have been performed in skeletal muscle exploring this pathway. However, recent observations from us and other laboratories have implicated UBR4, UBR5, and UBR7 in the regulation of the skeletal muscle proteome under growth and exercise stimuli (35–39). For instance, UBR5 and UBR4 have each been observed to be exercise responsive E3 ubiquitin ligases in human and mouse skeletal muscle under resistance and aerobic exercise models (35, 36, 38), with a change in phosphorylation status for UBR5 noted in multiple different exercise modalities (38). Further, UBR5 has been observed to increase in other models of mechanical loading (42) and UBR5 activity may be rapamycin insensitive (74) which compliments data collected in the current study in our Akt overexpression and UBR5 knockdown models. Our understanding of the transcriptional regulation of UBR5 and other UBR-related E3 ligases in skeletal muscle is limited. Previous literature has highlighted UBR5 DNA methylation status to coincide with changes in gene expression in human skeletal muscle under exercise training (35). Future studies are warranted to understand the transcriptional regulation of the UBR E3 ubiquitin ligases under different cellular conditions. To note, UBR5 has been reported to interact with p90RSK signaling via p90RSK phosphorylation sites on UBR5 (75), and we saw p90RSK activation with UBR5 suppression or increased UBR5 protein abundance in the presence of p90RSK phosphorylation, highlighting the potential responsiveness of UBR5 and UBR-box E3 ubiquitin ligases to changes in the protein synthetic machinery. These observations extend our understanding of E3 ubiquitin ligases in protein quality control and the regulation of skeletal muscle size.

Although very little is known about the substrates for the UBR-box E3 ubiquitin ligases, we observed an increase in K48-linked ubiquitinated proteins, total ubiquitinated protein levels and 26S proteasome activity under control and rapamycin diet conditions in the Akt overexpression model. Recent evidence by Kaiser and colleagues (19) has alluded to the proteasome system being controlled by mTORC1 via feedback inhibition of PKB/Akt. The authors utilized transgenic Akt-CA mice and TSC1 knockout models to manipulate mTORC1 activation and observed changes in proteasome biogenesis and activity as well as atrophy-related gene expression for specific E3 ubiquitin ligases in these models. It could be postulated that the increase in mTORC1-mediated translation results in a buildup of error containing peptides and misfolded proteins, which inadvertently increases the demand on the UPS for targeted protein degradation. These ideas warrant further investigation, and are highlighted in the literature drawing on observations reported using other cell and tissue models (76). In our current study, Akt overexpression and subsequent mTORC1 activation was induced for a shorter period of time compared to previous studies (15, 19, 20), and notably, we observed UPS-induction and markers of other protein quality control pathways (e.g. ER stress and autophagy) to be elevated under conditions where mTORC1 activity is low (e.g. rapamycin diet). Given, that it has previously been proposed that mTORC1 may control the UPS system and proteasome activity in skeletal muscle (19), the current study’s findings extend our observations for the UPS and protein quality control system being controlled under rapamycin-sensitive and insensitive signals. It is important to note that the chronic use of rapamycin can inhibit mTORC1 and mTORC2 (45). However, we observed increased phosphorylation of p90RSK and NDRG1 (mTORC2 substrate) with Akt overexpression and thus the activation of alternative protein synthetic pathways may require a response from a variety of protein quality control mechanisms such as the N-degron pathway in order to maintain proteome integrity.

Recent studies have alluded to the importance of the N-degron pathway being critical for protein quality control via its ability to recognize destabilized proteins that display a degron signal (31–33). Interestingly, with UBR5 knockdown, we observed an increase in protein abundance for UBR2, UBR4 and UBR7 which may highlight a compensatory response in order to maintain proteostasis due to increased protein synthesis in UBR5 KD transfected muscles. The response in UBR E3 ubiquitin ligases to skeletal muscle regrowth (21d to 60d) is notable given that all the UBR E3s measured in the current study were elevated in protein abundance as the tissue recovered following a period of inactivity. We recently hypothesized that the UBR E3 ubiquitin ligases work as a protein surveillance system under growth conditions in order to maintain proteome integrity during production of newly synthesized proteins (39). The observations from the current study are interesting because UBR-box E3 ligases all share the UBR-domain, but the mechanisms for ubiquitin transfer differs between RING, F-box and Hect E3 ubiquitin ligases which are all comprised in the UBR class of E3 ligases (24, 29, 77). The substrates for the UBR-box E3 ligases remain to be identified in vivo. Previous literature has suggested that different UBRs may elicit their function for ubiquitination under different cellular compartments (27, 31, 78). For instance, UBR1 and UBR2 are reported to target misfolded proteins for degradation in the cytosolic compartment (31) whereas the action of UBR5 might be dominant in the nucleus and regulating nuclear receptors and transcription factors (33, 79–81). However, these previously published observations remain to be confirmed in vivo and in the context of skeletal muscle plasticity.

## Limitations

The data reported within the current study is not without limitations. The use of in vivo electroporation allows for a targeted skeletal muscle approach to understand the molecular mechanisms of hypertrophy and atrophy (50). However, transfection efficiency can play a role in observations made at the biochemical level, as a “dilution” effect can occur whereby changes in gene/protein expression are diluted by a large proportion of non-transfected fibers, leading to an underestimation of the effect of the gene of interest (50). Our electroporation protocol has been optimized to maximize transfection efficiency whilst also limiting the level of potential muscle damage that might be observed (50, 51). Our current observations for the UBR-box E3 ligases in skeletal muscle remodeling are limited to only the tibialis anterior muscle, although we have performed similar electroporation experiments in the gastrocnemius muscle with the UBR5 RNAi construct and observed similar levels of muscle mass and fiber CSA decreases with UBR5 suppression (data not shown). In the current study, we observed changes in UBR-box E3 ligase protein abundance during the recovery of muscle mass following atrophy in the gastrocnemius complex muscles (Fig. 3), which might imply that their role is similar in all hindlimb skeletal muscles, but future studies are warranted. It should be noted that the rapamycin dose used in the current study is not without potential side effects at the whole body or physiological level with alterations in body weight and glucose metabolism being reported (82, 83). However, in the current study, we utilized the rapamycin diet intervention for a shorter period of time, compared to studies focused on rapamycin as an intervention for lifespan and healthspan (82, 83). Lastly, protein turnover and quality control are dynamic processes for maintaining proteostasis (76, 84, 85) and thus caution must be applied given that a single snapshot in time was measured for changes in protein quality control mechanisms when protein synthesis was genetically enhanced to induce skeletal muscle growth.

## Conclusions

The present study highlights a novel group of E3 ubiquitin ligases that respond to heightened protein synthesis under skeletal muscle growth or atrophy conditions via genetic manipulation or recovery following nerve crush injury. Little is known about the N-degron pathway in tissue health, and our initial observations provide important steps towards understanding this pathway. Future studies are warranted to identify protein substrates for these E3 ubiquitin ligases and explore their potential role in other tissues (e.g. heart, brain) given that they are ubiquitous E3 ligases. Overall, the UBR-box E3 ubiquitin ligases could be critical in the adaptive response to cellular stress where perturbations in proteostasis and tissue remodeling might occur in the context of aging, disease, and exercise.

## Data Availability

The data that support the findings of this study are available from the corresponding author upon reasonable request.

## Author Contributions

Conception and design of the experiments: SCB and DCH. Collection, analysis and interpretation of data: LMB, LGO, CAG, APS, DSW, SCB and DCH. Drafting the article or revising it critically for important intellectual content: LMB, LGO, CAG, APS, DSW, SCB and DCH. All authors read and approved the final version of the manuscript, and all authors listed qualify for authorship.

---

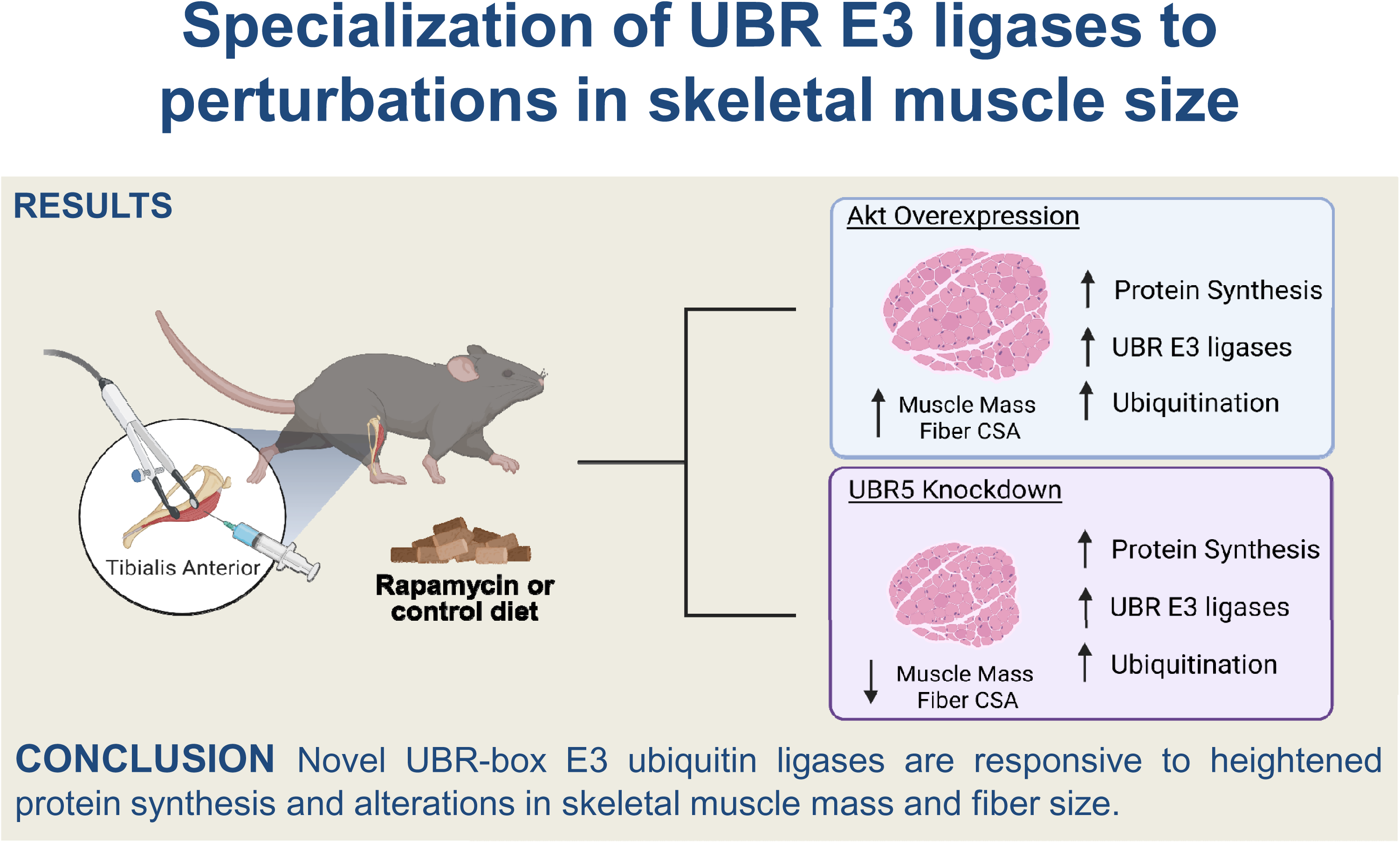

## References

1. Hughes DC, Ellefsen S, and Baar K. Adaptations to endurance and strength training. Cold Spring Harbor perspectives in medicine a029769, 2017.

2. Baehr LM, Hughes DC, Waddell DS, and Bodine SC. SnapShot: Skeletal muscle atrophy. Cell 185: 1618–1618. e1611, 2022.

3. Sharples AP, Hughes DC, Deane CS, Saini A, Selman C, and Stewart CE. Longevity and skeletal muscle mass: the role of IGF signalling, the sirtuins, dietary restriction and protein intake. Aging cell 14: 511–523, 2015.

4. Bodine SC, Stitt TN, Gonzalez M, Kline WO, Stover GL, Bauerlein R, Zlotchenko E, Scrimgeour A, Lawrence JC, and Glass DJ. Akt/mTOR pathway is a crucial regulator of skeletal muscle hypertrophy and can prevent muscle atrophy in vivo. Nature cell biology 3: 1014–1019, 2001.

5. Baar K, and Esser K. Phosphorylation of p70S6kcorrelates with increased skeletal muscle mass following resistance exercise. American Journal of Physiology-Cell Physiology 276: C120–C127, 1999.

6. West DW, Baehr LM, Marcotte GR, Chason CM, Tolento L, Gomes AV, Bodine SC, and Baar K. Acute resistance exercise activates rapamycin-sensitive and-insensitive mechanisms that control translational activity and capacity in skeletal muscle. The Journal of physiology 594: 453–468, 2016.

7. Lee Hamilton D, Philp A, MacKenzie MG, Patton A, Towler MC, Gallagher IJ, Bodine SC, and Baar K. Molecular brakes regulating mTORC1 activation in skeletal muscle following synergist ablation. American Journal of Physiology-Endocrinology and Metabolism 307: E365–E373, 2014.

8. You J-S, Dooley MS, Kim C-R, Kim E-J, Xu W, Goodman CA, and Hornberger TA. A DGKζ-FoxO-ubiquitin proteolytic axis controls fiber size during skeletal muscle remodeling. Science signaling 11: 2018.

9. Frey JW, Jacobs BL, Goodman CA, and Hornberger TA. A role for Raptor phosphorylation in the mechanical activation of mTOR signaling. Cellular Signalling 26: 313–322, 2014.

10. Goodman CA. Role of mTORC1 in mechanically induced increases in translation and skeletal muscle mass. Journal of applied physiology 127: 581–590, 2019.

11. Kim H-G, Guo B, and Nader GA. Regulation of ribosome biogenesis during skeletal muscle hypertrophy. Exercise and sport sciences reviews 47: 91–97, 2019.

12. Figueiredo VC, and McCarthy JJ. Regulation of ribosome biogenesis in skeletal muscle hypertrophy. Physiology 34: 30–42, 2019.

13. Wallace MA, Hughes DC, and Baar K. mTORC1 in the Control of Myogenesis and Adult Skeletal Muscle Mass. In: Molecules to Medicine with mTORElsevier, 2016, p. 37–56.

14. Castets P, Rion N, Théodore M, Falcetta D, Lin S, Reischl M, Wild F, Guérard L, Eickhorst C, Brockhoff M, Guridi M, Ibebunjo C, Cruz J, Sinnreich M, Rudolf R, Glass DJ, and Rüegg MA. mTORC1 and PKB/Akt control the muscle response to denervation by regulating autophagy and HDAC4. Nat Commun 10: 3187, 2019.

15. Marabita M, Baraldo M, Solagna F, Ceelen Judith Johanna M, Sartori R, Nolte H, Nemazanyy I, Pyronnet S, Kruger M, Pende M, and Blaauw B. S6K1 Is Required for Increasing Skeletal Muscle Force during Hypertrophy. Cell Reports 17: 501–513, 2016.

16. Bentzinger CF, Lin S, Romanino K, Castets P, Guridi M, Summermatter S, Handschin C, Tintignac LA, Hall MN, and Rüegg MA. Differential response of skeletal muscles to mTORC1 signaling during atrophy and hypertrophy. Skeletal muscle 3: 1–16, 2013.

17. Tang H, Inoki K, Brooks SV, Okazawa H, Lee M, Wang J, Kim M, Kennedy CL, Macpherson PC, and Ji X. mTORC1 underlies age-related muscle fiber damage and loss by inducing oxidative stress and catabolism. Aging cell 18: e12943, 2019.

18. Lai K-MV, Gonzalez M, Poueymirou WT, Kline WO, Na E, Zlotchenko E, Stitt TN, Economides AN, Yancopoulos GD, and Glass DJ. Conditional activation of akt in adult skeletal muscle induces rapid hypertrophy. Molecular and cellular biology 24: 9295–9304, 2004.

19. Kaiser MS, Milan G, Ham DJ, Lin S, Oliveri F, Chojnowska K, Tintignac LA, Mittal N, Zimmerli CE, and Glass DJ. Dual roles of mTORC1-dependent activation of the ubiquitin-proteasome system in muscle proteostasis. Communications biology 5: 1141, 2022.

20. Blaauw B, Canato M, Agatea L, Toniolo L, Mammucari C, Masiero E, Abraham R, Sandri M, Schiaffino S, and Reggiani C. Inducible activation of Akt increases skeletal muscle mass and force without satellite cell activation. The FASEB journal 23: 3896–3905, 2009.

21. Castets P, and Rüegg MA. MTORC1 determines autophagy through ULK1 regulation in skeletal muscle. Autophagy 9: 1435–1437, 2013.

22. Bodine SC, Latres E, Baumhueter S, Lai VK-M, Nunez L, Clarke BA, Poueymirou WT, Panaro FJ, Na E, and Dharmarajan K. Identification of ubiquitin ligases required for skeletal muscle atrophy. Science 294: 1704–1708, 2001.

23. Milan G, Romanello V, Pescatore F, Armani A, Paik J-H, Frasson L, Seydel A, Zhao J, Abraham R, and Goldberg AL. Regulation of autophagy and the ubiquitin–proteasome system by the FoxO transcriptional network during muscle atrophy. Nature communications 6: 6670, 2015.

24. Hughes DC, Goodman CA, Baehr LM, Gregorevic P, and Bodine SC. A critical discussion on the relationship between E3 ubiquitin ligases, protein degradation, and skeletal muscle wasting: it’s not that simple. American Journal of Physiology-Cell Physiology 325: C1567–C1582, 2023.

25. Varshavsky A. N-degron and C-degron pathways of protein degradation. Proceedings of the National Academy of Sciences 116: 358–366, 2019.

26. Varshavsky A. The N-end rule pathway and regulation by proteolysis. Protein science 20: 1298–1345, 2011.

27. Sherpa D, Chrustowicz J, and Schulman BA. How the ends signal the end: Regulation by E3 ubiquitin ligases recognizing protein termini. Molecular Cell 82: 1424–1438, 2022.

28. Choi WS, Jeong B-C, Joo YJ, Lee M-R, Kim J, Eck MJ, and Song HK. Structural basis for the recognition of N-end rule substrates by the UBR box of ubiquitin ligases. Nature structural & molecular biology 17: 1175–1181, 2010.

29. Tasaki T, Mulder LC, Iwamatsu A, Lee MJ, Davydov IV, Varshavsky A, Muesing M, and Kwon YT. A family of mammalian E3 ubiquitin ligases that contain the UBR box motif and recognize N-degrons. Molecular and cellular biology 2005.

30. Hunt LC, Schadeberg B, Stover J, Haugen B, Pagala V, Wang Y-D, Puglise J, Barton ER, Peng J, and Demontis F. Antagonistic control of myofiber size and muscle protein quality control by the ubiquitin ligase UBR4 during aging. Nature Communications 12: 1418, 2021.

31. Nillegoda NB, Theodoraki MA, Mandal AK, Mayo KJ, Ren HY, Sultana R, Wu K, Johnson J, Cyr DM, and Caplan AJ. Ubr1 and Ubr2 function in a quality control pathway for degradation of unfolded cytosolic proteins. Molecular biology of the cell 21: 2102–2116, 2010.

32. Shimshon A, Dahan K, Israel-Gueta M, Olmayev-Yaakobov D, Timms RT, Bekturova A, Makaros Y, Elledge SJ, and Koren I. Dipeptidyl peptidases and E3 ligases of N-degron pathways cooperate to regulate protein stability. Journal of Cell Biology 223: e202311035, 2024.

33. Tsai JM, Aguirre JD, Li Y-D, Brown J, Focht V, Kater L, Kempf G, Sandoval B, Schmitt S, and Rutter JC. UBR5 forms ligand-dependent complexes on chromatin to regulate nuclear hormone receptor stability. Molecular cell 83: 2753–2767. e2710, 2023.

34. Solomon V, Lecker SH, and Goldberg AL. The N-end rule pathway catalyzes a major fraction of the protein degradation in skeletal muscle. Journal of Biological Chemistry 273: 25216–25222, 1998.

35. Seaborne RA, Strauss J, Cocks M, Shepherd S, O’Brien TD, van Someren KA, Bell PG, Murgatroyd C, Morton JP, Stewart CE, and Sharples AP. Human Skeletal Muscle Possesses an Epigenetic Memory of Hypertrophy. Sci Rep 8: 1898, 2018.

36. Chambers TL, Dimet-Wiley A, Keeble AR, Haghani A, Lo WJ, Kang G, Brooke R, Horvath S, Fry CS, and Watowich SJ. Methylome–proteome integration after late-life voluntary exercise training reveals regulation and target information for improved skeletal muscle health. The Journal of Physiology 603: 211–237, 2025.

37. Roberts MD, Ruple BA, Godwin JS, McIntosh MC, Chen S-Y, Kontos NJ, Agyin-Birikorang A, Michel M, Plotkin DL, and Mattingly ML. A novel deep proteomic approach in human skeletal muscle unveils distinct molecular signatures affected by aging and resistance training. Aging (Albany NY) 16: 6631, 2024.

38. Blazev R, Carl CS, Ng Y-K, Molendijk J, Voldstedlund CT, Zhao Y, Xiao D, Kueh AJ, Miotto PM, and Haynes VR. Phosphoproteomics of three exercise modalities identifies canonical signaling and C18ORF25 as an AMPK substrate regulating skeletal muscle function. Cell Metabolism 34: 1561–1577. e1569, 2022.

39. Hughes DC, and Bodine SC. UBR5: A New Player in Protein Quality Control for Skeletal Muscle Growth and Remodeling. Exercise and Sport Sciences Reviews 53: 205–213, 2025.

40. Gao S, Zhang G, Zhang Z, Zhu JZ, Li L, Zhou Y, Rodney GG, Jr., Abo-Zahrah RS, Anderson L, Garcia JM, Kwon YT, and Li YP. UBR2 targets myosin heavy chain IIb and IIx for degradation: Molecular mechanism essential for cancer-induced muscle wasting. Proc Natl Acad Sci U S A 119: e2200215119, 2022.

41. Kwak KS, Zhou X, Solomon V, Baracos VE, Davis J, Bannon AW, Boyle WJ, Lacey DL, and Han HQ. Regulation of protein catabolism by muscle-specific and cytokine-inducible ubiquitin ligase E3alpha-II during cancer cachexia. Cancer Res 64: 8193–8198, 2004.

42. Seaborne RA, Hughes DC, Turner DC, Owens DJ, Baehr LM, Gorski P, Semenova EA, Borisov OV, Larin AK, Popov DV, Generozov EV, Sutherland H, Ahmetov II, Jarvis JC, Bodine SC, and Sharples AP. UBR5 is a novel E3 ubiquitin ligase involved in skeletal muscle hypertrophy and recovery from atrophy. The Journal of Physiology 597: 3727–3749, 2019.

43. Hughes DC, Turner DC, Baehr LM, Seaborne RA, Viggars M, Jarvis JC, Gorski PP, Stewart CE, Owens DJ, Bodine SC, and Sharples AP. Knockdown of the E3 Ubiquitin ligase UBR5 and its role in skeletal muscle anabolism. American Journal of Physiology-Cell Physiology 320: C45–C56, 2021.

44. Hunt LC, Stover J, Haugen B, Shaw TI, Li Y, Pagala VR, Finkelstein D, Barton ER, Fan Y, and Labelle M. A key role for the ubiquitin ligase UBR4 in myofiber hypertrophy in Drosophila and mice. Cell reports 28: 1268–1281. e1266, 2019.

45. Sarbassov DD, Ali SM, Sengupta S, Sheen JH, Hsu PP, Bagley AF, Markhard AL, and Sabatini DM. Prolonged rapamycin treatment inhibits mTORC2 assembly and Akt/PKB. Mol Cell 22: 159–168, 2006.

46. Ham DJ, Börsch A, Chojnowska K, Lin S, Leuchtman AB, Ham AS, Thürkauf M, Delezie J, Furrer R, Burri D, Sinnreich M, Handschin C, Tintignac LA, Zavolan M, Mittal N, and Rüegg MA. Distinct and additive effects of calorie restriction and rapamycin in aging skeletal muscle. Nature Communications 13: 2025, 2022.

47. Ham DJ, Börsch A, Lin S, Thürkauf M, Weihrauch M, Reinhard JR, Delezie J, Battilana F, Wang X, and Kaiser MS. The neuromuscular junction is a focal point of mTORC1 signaling in sarcopenia. Nature Communications 11: 1–21, 2020.

48. Chiao YA, Kolwicz SC, Basisty N, Gagnidze A, Zhang J, Gu H, Djukovic D, Beyer RP, Raftery D, and MacCoss M. Rapamycin transiently induces mitochondrial remodeling to reprogram energy metabolism in old hearts. Aging (Albany NY) 8: 314, 2016.

49. Ebert SM, Dyle MC, Kunkel SD, Bullard SA, Bongers KS, Fox DK, Dierdorff JM, Foster ED, and Adams CM. Stress-induced skeletal muscle Gadd45a expression reprograms myonuclei and causes muscle atrophy. J Biol Chem 287: 27290–27301, 2012.

50. Hughes DC, Hardee JP, Waddell DS, and Goodman CA. CORP: Gene delivery into murine skeletal muscle using in vivo electroporation. Journal of Applied Physiology 133: 41–59, 2022.

51. Hughes DC, Baehr LM, Driscoll JR, Lynch SA, Waddell DS, and Bodine SC. Identification and characterization of Fbxl22, a novel skeletal muscle atrophy-promoting E3 ubiquitin ligase. Am J Physiol Cell Physiol 319: C700–c719, 2020.

52. Baehr LM, Hughes DC, Lynch SA, Van Haver D, Maia TM, Marshall AG, Radoshevich L, Impens F, Waddell DS, and Bodine SC. Identification of the MuRF1 skeletal muscle ubiquitylome through quantitative proteomics. Function 2: zqab029, 2021.

53. Wen Y, Murach KA, Vechetti IJ, Jr., Fry CS, Vickery C, Peterson CA, McCarthy JJ, and Campbell KS. MyoVision: software for automated high-content analysis of skeletal muscle immunohistochemistry. J Appl Physiol (1985) 124: 40–51, 2018.

54. Viggars MR, Wen Y, Peterson CA, and Jarvis JC. Automated cross-sectional analysis of trained, severely atrophied, and recovering rat skeletal muscles using MyoVision 2.0. J Appl Physiol (1985) 132: 593–610, 2022.

55. Parlee SD, Lentz SI, Mori H, and MacDougald OA. Quantifying size and number of adipocytes in adipose tissue. Methods Enzymol 537: 93–122, 2014.

56. Castets P, Lin S, Rion N, Di Fulvio S, Romanino K, Guridi M, Frank S, Tintignac LA, Sinnreich M, and Rüegg MA. Sustained activation of mTORC1 in skeletal muscle inhibits constitutive and starvation-induced autophagy and causes a severe, late-onset myopathy. Cell Metab 17: 731–744, 2013.

57. Ching JK, Elizabeth SV, Ju JS, Lusk C, Pittman SK, and Weihl CC. mTOR dysfunction contributes to vacuolar pathology and weakness in valosin-containing protein associated inclusion body myopathy. Hum Mol Genet 22: 1167–1179, 2013.

58. Goodman CA, and Hornberger TA. Measuring protein synthesis with SUnSET: a valid alternative to traditional techniques? Exerc Sport Sci Rev 41: 107–115, 2013.

59. Baehr LM, Tunzi M, and Bodine SC. Muscle hypertrophy is associated with increases in proteasome activity that is independent of MuRF1 and MAFbx expression. Front Physiol 5: 69, 2014.

60. Gomes AV, Waddell DS, Siu R, Stein M, Dewey S, Furlow JD, and Bodine SC. Upregulation of proteasome activity in muscle RING finger 1-null mice following denervation. The FASEB Journal 26: 2986–2999, 2012.

61. Heinemeier KM, Olesen JL, Haddad F, Schjerling P, Baldwin KM, and Kjaer M. Effect of unloading followed by reloading on expression of collagen and related growth factors in rat tendon and muscle. Journal of applied physiology 106: 178–186, 2009.

62. Baehr LM, West DWD, Marcotte G, Marshall AG, De Sousa LG, Baar K, and Bodine SC. Age-related deficits in skeletal muscle recovery following disuse are associated with neuromuscular junction instability and ER stress, not impaired protein synthesis. Aging 8: 127–146, 2016.

63. Baehr LM, West DWD, Marshall AG, Marcotte GR, Baar K, and Bodine SC. Muscle-specific and age-related changes in protein synthesis and protein degradation in response to hindlimb unloading in rats. J Appl Physiol (1985) 122: 1336–1350, 2017.

64. Hughes DC, Marcotte GR, Baehr LM, West DW, Marshall AG, Ebert SM, Davidyan A, Adams CM, Bodine SC, and Baar K. Alterations in the muscle force transfer apparatus in aged rats during unloading and reloading: impact of microRNA-31. The Journal of physiology 596: 2883–2900, 2018.

65. Biever A, Puighermanal E, Nishi A, David A, Panciatici C, Longueville S, Xirodimas D, Gangarossa G, Meyuhas O, Hervé D, Girault J-A, and Valjent E. PKA-Dependent Phosphorylation of Ribosomal Protein S6 Does Not Correlate with Translation Efficiency in Striatonigral and Striatopallidal Medium-Sized Spiny Neurons. The Journal of Neuroscience 35: 4113–4130, 2015.

66. Bonito-Oliva A, Pallottino S, Bertran-Gonzalez J, Girault J-A, Valjent E, and Fisone G. Haloperidol promotes mTORC1-dependent phosphorylation of ribosomal protein S6 via dopamine– and cAMP-regulated phosphoprotein of 32 kDa and inhibition of protein phosphatase-1. Neuropharmacology 72: 197–203, 2013.

67. Roux PP, Shahbazian D, Vu H, Holz MK, Cohen MS, Taunton J, Sonenberg N, and Blenis J. RAS/ERK signaling promotes site-specific ribosomal protein S6 phosphorylation via RSK and stimulates cap-dependent translation. J Biol Chem 282: 14056–14064, 2007.

68. Sartori R, Romanello V, and Sandri M. Mechanisms of muscle atrophy and hypertrophy: Implications in health and disease. Nature Communications 12: 1–12, 2021.

69. Ogasawara R, and Suginohara T. Rapamycin-insensitive mechanistic target of rapamycin regulates basal and resistance exercise-induced muscle protein synthesis. The FASEB Journal 32: 5824–5834, 2018.

70. Ogasawara R, Knudsen JR, Li J, Ato S, and Jensen TE. Rapamycin and mTORC2 inhibition synergistically reduce contraction-stimulated muscle protein synthesis. The Journal of physiology 598: 5453–5466, 2020.

71. You J-S, McNally RM, Jacobs BL, Privett RE, Gundermann DM, Lin K-H, Steinert ND, Goodman CA, and Hornberger TA. The role of raptor in the mechanical load-induced regulation of mTOR signaling, protein synthesis, and skeletal muscle hypertrophy. The FASEB Journal 33: 4021, 2018.

72. Burd NA, Holwerda AM, Selby KC, West DW, Staples AW, Cain NE, Cashaback JG, Potvin JR, Baker SK, and Phillips SM. Resistance exercise volume affects myofibrillar protein synthesis and anabolic signalling molecule phosphorylation in young men. The Journal of physiology 588: 3119–3130, 2010.

73. Williamson D, Gallagher P, Harber M, Hollon C, and Trappe S. Mitogen-activated protein kinase (MAPK) pathway activation: effects of age and acute exercise on human skeletal muscle. The Journal of physiology 547: 977–987, 2003.

74. Steinert ND, Potts GK, Wilson GM, Klamen AM, Lin KH, Hermanson JB, McNally RM, Coon JJ, and Hornberger TA. Mapping of the contraction-induced phosphoproteome identifies TRIM28 as a significant regulator of skeletal muscle size and function. Cell Rep 34: 108796, 2021.

75. Cho JH, Kim SA, Seo Y-S, Park SG, Park BC, Kim J-H, and Kim S. The p90 ribosomal S6 kinase– UBR5 pathway controls Toll-like receptor signaling via miRNA-induced translational inhibition of tumor necrosis factor receptor–associated factor 3. Journal of Biological Chemistry 292: 11804–11814, 2017.

76. Pla-Prats C, and Thomä NH. Quality control of protein complex assembly by the ubiquitin– proteasome system. Trends in cell biology 32: 696–706, 2022.

77. Komander D, and Rape M. The Ubiquitin Code. Annual Review of Biochemistry 81: 203–229, 2012.

78. Hogan AK, Sathyan KM, Willis AB, Khurana S, Srivastava S, Zasadzińska E, Lee AS, Bailey AO, Gaynes MN, and Huang J. UBR7 acts as a histone chaperone for post-nucleosomal histone H3. The EMBO journal 40: e108307, 2021.

79. Taherbhoy AM, and Daniels DL. Harnessing UBR5 for targeted protein degradation of key transcriptional regulators. Trends in Pharmacological Sciences 2023.

80. Duan C-y, Li Y, Zhi H-y, Tian Y, Huang Z-y, Chen S-P, Zhang Y, Liu Q, Zhou L, and Jiang X-g. E3 ubiquitin ligase UBR5 modulates circadian rhythm by facilitating the ubiquitination and degradation of the key clock transcription factor BMAL1. Acta Pharmacologica Sinica 45: 1793–1808, 2024.

81. Mark KG, Kolla S, Aguirre JD, Garshott DM, Schmitt S, Haakonsen DL, Xu C, Kater L, Kempf G, and Martínez-González B. Orphan quality control shapes network dynamics and gene expression. Cell 186: 3460–3475. e3423, 2023.

82. Baghdadi M, Nespital T, Monzó C, Deelen J, Grönke S, and Partridge L. Intermittent rapamycin feeding recapitulates some effects of continuous treatment while maintaining lifespan extension. Mol Metab 81: 101902, 2024.

83. Miller RA, Harrison DE, Astle CM, Fernandez E, Flurkey K, Han M, Javors MA, Li X, Nadon NL, Nelson JF, Pletcher S, Salmon AB, Sharp ZD, Van Roekel S, Winkleman L, and Strong R. Rapamycin-mediated lifespan increase in mice is dose and sex dependent and metabolically distinct from dietary restriction. Aging Cell 13: 468–477, 2014.

84. O’Reilly CL, Bodine SC, and Miller BF. Current limitations and future opportunities of tracer studies of muscle ageing. The Journal of Physiology 603: 7, 2023.

85. Brandman O, Stewart-Ornstein J, Wong D, Larson A, Williams CC, Li G-W, Zhou S, King D, Shen PS, and Weibezahn J. A ribosome-bound quality control complex triggers degradation of nascent peptides and signals translation stress. Cell 151: 1042–1054, 2012.

